# Multiple behavioural mechanisms shape development in a highly social cichlid fish

**DOI:** 10.1101/2023.04.14.536957

**Authors:** Isabela P. Harmon, Emily A. McCabe, Madeleine R. Vergun, Julia Weinstein, Hannah L. Graves, Deijah D. Bradley, Clare M. Boldt, June Lee, Jessica M. Maurice, Tessa K. Solomon-Lane

**Affiliations:** Scripps College, Claremont, CA; Claremont McKenna College, Claremont, CA; Pitzer College, Claremont, CA

**Keywords:** Behavioural syndrome, Cortisol, Early-life experience, Early-life social effects, Ontogeny, Hypothalamic-pituitary-adrenal axis

## Abstract

Early-life social experiences shape adult phenotype, yet the underlying behavioural mechanisms remain poorly understood. We manipulated early-life social experience in the highly social African cichlid fish *Astatotilapia burtoni* to investigate the effects on behaviour and neuroendocrine stress axis function. Juveniles experienced different numbers of early-life social partners in stable pairs (1 partner), stable groups (6 fish; 5 partners), and socialized pairs (a novel fish was exchanged every 5 days; 5 partners). Treatments differed in group size (groups vs. pairs) and stability (stable vs. socialized). We then measured behaviour in multiple contexts and collected water-borne cortisol. We found effects of treatment on behaviour across all assays: open field exploration, social cue investigation, dominant behaviour, and subordinate behaviour. Cortisol did not differ across treatments. Principal components (PC) analysis revealed robust co- variation of behaviour across contexts, including with cortisol, to form behavioural syndromes sensitive to early-life social experience. PC1 (25.1%) differed by numbers of social partners: juveniles with more social partners were more active during the social cue investigation, spent less time in the territory, and were more interactive as dominants. Differences in PC5 (8.5%) were based on stability: socialized pairs were more dominant, spent less time in and around the territory, were more socially investigative, and had lower cortisol than stable groups or pairs. Behaviour observations in the home tanks provided further insights into the behavioural mechanisms underlying these effects. These results contribute to our understanding of how early-life social experiences are accrued and exert strong, lasting effects on adult phenotype.

## INTRODUCTION

Early-life social environments and experiences are potent drivers of developmental plasticity for social species and, as a result, can have strong, long-term effects on organismal phenotype (Bateson, 2001; Bateson et al., 2004; Kuijper & Johnstone, 2019; Taborsky, 2017; Weaver, 2009). Early-life social effects have been documented across vertebrate taxa (e.g., Arnold & Taborsky, 2010; Bölting & von Engelhardt, 2017; Champagne & Curley, 2005; Moretz et al., 2007; Perkeybile & Bales, 2017; White et al., 2010), yet the specific attributes of early social environments and experiences that cause phenotypic changes are often unidentified (Kasumovic, 2013; Taborsky, 2016). Social interactions and stimuli can make up a substantial part of juvenile experience (Kohn, 2019). Depending on the species and group structure, early social interactions can involve parents (maternal, paternal, or biparental care) (Champagne & Curley, 2005; McClelland et al., 2011; Perkeybile et al., 2013), parental helpers (Arnold & Taborsky, 2010; Taborsky et al., 2012), siblings (Branchi et al., 2013; Buist et al., 2013; D’Andrea et al., 2007; Monclús et al., 2012), peers (Ahloy Dallaire & Mason, 2017; Bölting & von Engelhardt, 2017; Förster & Cords, 2005; Moretz et al., 2007; Weinstein et al., 2014), and other members of the group (Bray, Murray, et al., 2021; Förster & Cords, 2005; Jin et al., 2015), as well as observations of others interacting (Clay & de Waal, 2013; Desjardins et al., 2012; Oliveira et al., 1998). Identifying the specific, proximate causes—the behavioural mechanisms— is critical to understanding how gene-by-environment interactions shape processes of developmental plasticity and behavioural developmental trajectories. Given the fitness and health consequences of social behaviour (Bennett et al., 2006; Meyer-Lindenberg & Tost, 2012; Silk, 2007; Solomon-Lane et al., 2015; Wilson, 1980), the developmental origins of adult behavioural phenotype are particularly important to understand, including as a target for natural selection.

Manipulating the early social environment is a common approach to studying early-life social effects. For example, rearing animals in groups of different sizes and/or complexities often has significant effects on development and phenotype (reviewed in Taborsky, 2016). In ravens (*Corvus corax*), family size affected social attentiveness (Gallego-Abenza et al., 2022); in zebra finches (*Taeniopygia guttata*), rearing group size and age diversity of early-life flocks affected courtship and aggressive behaviour, as well as plumage development (Bölting & von Engelhardt, 2017); in mice, communal nesting affected social skills and neuroendocrine function, with separate effects of maternal care and peer interactions (Branchi et al., 2009, 2013); and in Daffodil cichlids (*Neolamprologus pulcher*), the presence of parents and brood helpers increased social competence (Arnold & Taborsky, 2010; Taborsky et al., 2012). In general, larger social groups are more complex than smaller groups, and early exposure to social complexity tends to benefit social skills and competence (e.g., Branchi et al., 2009, 2013; Fischer et al., 2015; White et al., 2010). However, individuals accrue different social experiences, even within shared environments. For example, mouse pups in mixed-age, communal nests interact with siblings at varying rates and receive different levels of maternal care (Branchi et al., 2013); infant blue monkeys (*Cercopithecus mitis stuhlmanni*) receive varying rates of allomaternal care, and from different non-mothers in the group (Förster & Cords, 2005); immature male chimpanzees socially associate with adult males at different rates (Bray, Feldblum, et al., 2021); and young male long-tailed manakins occupy varied positions in the social network (McDonald, 2007). This variation can have long-term effects on social decision-making and behaviour (e.g., Branchi et al., 2013; Bray, Murray, et al., 2021; McDonald, 2007). Therefore, to identify the behavioural mechanisms underlying behavioural development, it is necessary to observe individuals in the rearing environment and test mechanistic hypotheses directly by manipulating the quality and/or quantity of social experiences (Kasumovic, 2013; Taborsky, 2016).

Early social experiences exert long-term effects on organismal phenotype through persistent changes in underlying neural and neuroendocrine mechanisms (e.g., Antunes et al., 2021; Branchi et al., 2013; Champagne & Curley, 2005; McClelland et al., 2011). The neuroendocrine stress axis, or hypothalamic-pituitary-adrenal (interrenal in fish; HPA/I) axis, is a highly-conserved mechanism underlying early-life effects, including social effects (Banerjee et al., 2012; Champagne & Curley, 2005; Crespi & Denver, 2005; Ensminger et al., 2018; Francis et al., 1999; Jonsson & Jonsson, 2014; McClelland et al., 2011; Taborsky, 2016). The HPA/I axis translates environmental conditions and experiences into coordinated physiological and behavioural responses through a neuroendocrine cascade that initiates in response to an environmental stressor. Detection of a stressor, which can include any external condition that disrupts or threatens to disrupt homeostasis, leads to the release of corticotropin-releasing factor (CRF) from the hypothalamus. The pituitary responds to CRF with the release of adrenocorticotropic (ACTH) hormone, which signals for the adrenal (or interrenal) glands to release glucocorticoids (cortisol in fishes) into circulation (Denver, 2009; Lowry & Moore, 2006; Wendelaar Bonga, 1997). Early-life social experiences shape HPA/I axis function in multiple ways (Champagne & Curley, 2005; Francis et al., 1999; McClelland et al., 2011; Spencer, 2017; Taborsky, 2016; Turecki & Meaney, 2016). For example, peer-reared rhesus macaques (*Macaca mulatta*) had higher baseline and stress-induced levels of ACTH and cortisol, as well as a more reactive HPA axis, compared to maternal-reared juveniles (Stevens et al., 2009). Similarly, zebra finch chicks reared with only their fathers had hyperresponsive HPA axes, along with altered levels of neural glucocorticoid receptor (GR) and mineralocorticoid receptor (MR) expression, compared to biparental rearing (Banerjee et al., 2012). In *N. pulcher* cichlids, rearing with or without parents affected *gr1* expression in the telencephalon (Nyman et al., 2018), and rearing with or without brood helpers altered neural *crf* and *gr* expression, as well as the ratio of *mr* to *gr1* (Taborsky et al., 2013). The HPA/I axis is also an important source of individual variation in social behaviour (Boogert et al., 2014; Dettmer et al., 2017; Farine et al., 2015; Freeman et al., 2021; Pryce et al., 2011; Reyes-Contreras et al., 2019; Sih, 2011) and health (e.g., Turecki & Meaney, 2016).

In this study, we used Burton’s Mouthbrooder (*Astatotilapia burtoni*), a highly social cichlid fish and model system in social neuroscience (Fernald & Maruska, 2012; Hofmann, 2003; Stevenson et al., 2018), to investigate the behavioural mechanisms through which early-life social experiences affect behaviour and HPI axis function. The vast majority of research on this species has focused on adults, which form mixed-sex, hierarchical social communities. Dominant males are colourful, territorial, and reproductively active. In contrast, subordinate males are drab in coloration, shoal with females, and are reproductively suppressed. Social status is socially-regulated, and males regularly transition between dominant and subordinate positions (Fernald & Maruska, 2012; Hofmann, 2003). Juveniles also form status relationships (Solomon-Lane et al., 2022), and both juveniles and adults express a suite of highly-conserved social behaviours (Fernald & Hirata, 1979; Fernald & Maruska, 2012; Solomon-Lane et al., 2022; Weitekamp & Hofmann, 2017). Although sex and social status have strong effects on adult behaviour, individual variation persists, including in the frequency and quality of behavioural expression, such as aggression, territoriality, courtship, cooperation, reproductive behaviour, and maternal behaviour (Alward et al., 2021; Friesen et al., 2022; Fulmer et al., 2017; Kidd et al., 2013; Maruska, Becker, et al., 2013; Renn et al., 2009; Weitekamp & Hofmann, 2017); tenure in a dominant vs. subordinate role (Hofmann et al., 1999); female mate choice (Kidd et al., 2013); social learning (Rodriguez-Santiago et al., 2020); and cognition (Wallace & Hofmann, 2021). HPI axis function also varies among adults (Alcazar et al., 2016; Chen & Fernald, 2008; Clement et al., 2005; Dijkstra et al., 2017; Greenwood et al., 2003; Korzan et al., 2014; Maruska et al., 2022) and is highly responsive to social context, such as social status and changes in status (Carpenter et al., 2014; Chen & Fernald, 2008; Clement et al., 2005; Fox et al., 1997; Huffman et al., 2015; Korzan et al., 2014; Maruska et al., 2022; Maruska, Zhang, et al., 2013; Parikh et al., 2006), group social dynamics (Maguire et al., 2021), social habituation (Weitekamp & Hofmann, 2017), and in response to an intruder (Alcazar et al., 2016; Weitekamp et al., 2017).

Developmental plasticity may be an important source of individual variation in behaviour and HPI axis function among adults (Fernald & Hirata, 1979; Fraley & Fernald, 1982; Solomon-Lane & Hofmann, 2019). We have previously demonstrated early-life social effects in juvenile *A. burtoni* in response to early-life social group size. Juveniles reared in social groups (16 fish) developed to be more active, more socially interactive, and less submissive in a subordinate role compared to pair-reared juveniles. Whole brain gene expression related to HPI axis function was also altered, including elevated GR1a expression in group-reared animals and tight co-expression of candidate genes (including GR1a, GR1b, GR2, MR, and CRF) in pair-reared, but not group-reared or isolated, juveniles (Solomon-Lane & Hofmann, 2019). Juveniles in the previous study were not observed in the rearing environment, and there are multiple, possible behavioural mechanisms responsible for these effects (Solomon-Lane & Hofmann, 2019; Taborsky, 2016). Individuals reared in larger and/or more complex groups may interact socially at higher rates, experience a greater diversity of types of social interaction (e.g., affiliative, aggressive, cooperative, etc.), gain experience in multiple social roles (e.g., dominant and subordinate, younger and older) (Chase et al., 2022; Solomon-Lane et al., 2022; Williamson et al., 2016), have more social partners, have social partners of varied identities / traits (e.g., age, body size, life history stage diversity) (Arnold & Taborsky, 2010; Bölting & von Engelhardt, 2017; Branchi & Alleva, 2006; Taborsky et al., 2012; White et al., 2010), experience interactions involving more actors simultaneously (i.e., 3-way, or more, interactions), and learn indirectly by watching others (e.g., Desjardins et al., 2012; Oliveira et al., 1998)). For juvenile *A. burtoni*, group size and the relative body sizes of individuals affects social behaviour and group structure. For example, pairs and triads containing same-sized fish are less hierarchical (Solomon-Lane et al., 2022).

In this study, we tested the hypothesis that the number of social partners experienced by juvenile *A. burtoni* during early-life is a behavioural mechanism that shapes behavioural development and HPI axis signalling. The number of social partners is one clear way that groups and pairs differ (Solomon-Lane & Hofmann, 2019). There is also evidence from other species that this may be a meaningful attribute of social experience that varies across individuals. Assortative social interactions have been observed in a variety of species. For example, more socially interactive juvenile yellow-bellied marmots (*Marmota flaviventris*) had more novel social partners (Monclús et al., 2012); juvenile male geladas (*Theropithecus gelada*) had more novel play partners than juvenile females (Barale et al., 2015); juvenile rhesus macaques exhibit variation in the number of friendships initiated and reciprocated (Weinstein et al., 2014); and bold three-spined sticklebacks (*Gasterosteus aculeatus*) had a larger number of social contacts compared to shy fish (Pike et al., 2008). For juvenile *A. burtoni* in pairs and triads, individuals are not equally likely to initiate interactions (Solomon-Lane et al., 2022), suggesting that in larger, more naturalistic groups, social network connectivity may vary considerably, as it does in adults (Maguire et al., 2021). Direct manipulations of social partner numbers can also affect social behavioural development. For example, juvenile male brown-headed cowbirds reared in dynamic flocks outcompeted males reared in stable flocks of the same size (White et al., 2010).

To manipulate the number of social partners, we reared juveniles in stable pairs (2 fish, 1 partner each), in stable groups (6 fish, 5 partners each), or in socialized pairs, in which one member of the pair was changed 5 times over the course of the ∼1 month experiment (2 fish at a time in the pair, 5 total partners each) (Figure 1). This design also allowed us to compare the effects of group size (pair vs. group) and social stability (stable vs. socialized). We observed social interactions in the different rearing environments to quantify social experience and then tested the effects on individual social behaviour through a series of behaviour assays. We hypothesized that the number of social partners would be the strongest influence on behavioural phenotype. If the number of early-life social partners was an important behavioural mechanism underlying group- vs. pair-reared development in Solomon-Lane & Hofmann (2019), then in this study, we predict that juveniles from the stable groups and socialized pairs will be more active and socially interactive in novel social context, as well as less submissive in a subordinate role. To test the hypothesis that early-life social experience affects HPI axis signalling, we collected water-borne cortisol following the individual behaviour assays. By investigating specific behavioural mechanisms underlying behavioural development and neuroendocrine function, this research can provide insights into developmental plasticity processes and uncover the origins of adult phenotype.

**Figure 1:**
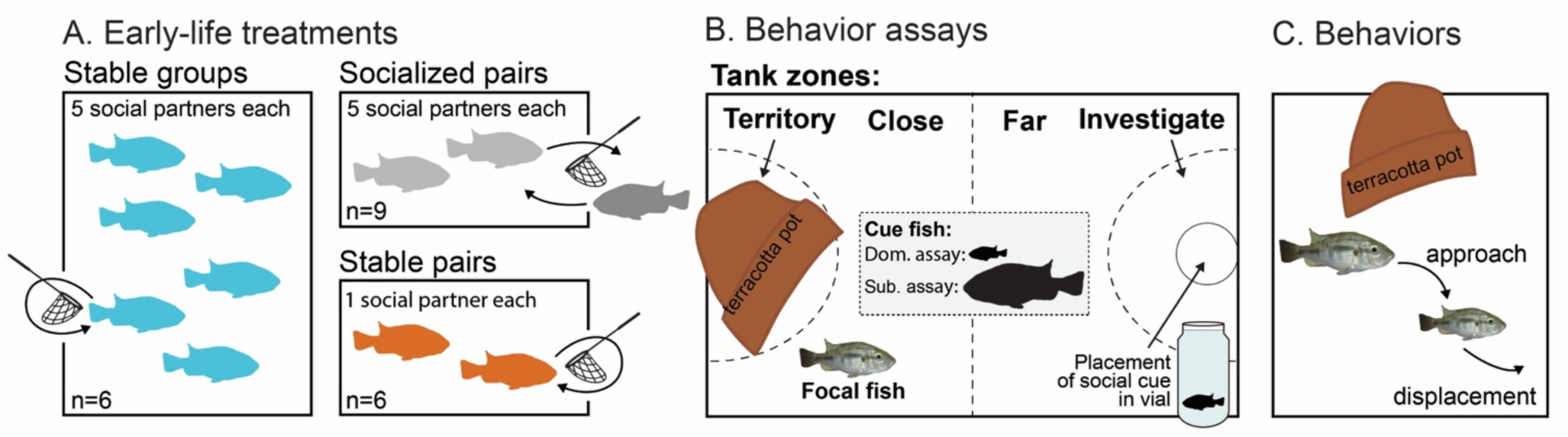
A) Juvenile fish were reared in stable groups (n=6, 6 fish each), stable pairs (n=6, 2 fish each), or socialized pairs (n=9, 2 fish each). Every 5 days in the socialized pairs, one fish of the pair was removed, and a novel juvenile was introduced. The novel juvenile came from a different socialized pair, and the removed juvenile became the novel partner for a socialized pair. This was repeated 5 times so that socialized fish had a total of 5 social partners – equal to the stable group fish. One fish per stable group and pair was also removed from their tank with a hand net as a control. This fish was immediately returned to its home tank. B) After 26 days in these rearing environments, individual juvenile behaviour was quantified in a novel experimental tank. The tank contained a terracotta pot shard, and black lines (drawn in permanent marker) divided the tank into four zones: territory, close, far, and investigate. In the open field exploration, the focal fish was alone in the tank. In the social cue investigation, a small juvenile cue fish inside of a scintillation vial was placed in the circle in the investigate zone. In the dominance assay, a freely-swimming novel cue fish (smaller than the focal) was added to the tank. In the subordinate assay, a freely-swimming novel cue fish (larger than the focal) was added to the tank. C) In the dominance and subordinate assay, we quantified the number of approaches, where one fish swims within three body lengths directly towards any part of another fish. If the approached fish swam away in any direction, it was counted as a displacement.

## METHODS

### Animals

The juvenile *A. burtoni* used in this experiment were bred in the laboratory. The breeding adults are a laboratory population that descended ∼65-70 generations from a wild caught stock from Lake Tanganyika (Fernald & Hirata, 1977). The adults are housed in naturalistic, mixed-sex breeding communities. Dominant males court females to lay eggs in his territory, after which she immediately scoops up the eggs into her mouth, where the male fertilizes them. The developing larvae are brooded in the female’s buccal cavity for ∼14 days before being released. In the wild, and under some laboratory conditions, females display maternal behaviour by guarding their offspring for 10 days or more after initially releasing the free-swimming fry from her mouth (Renn et al., 2009).

In this study, 86 fry were removed from the buccal cavities of 7 mothers approximately 5-7 days after fertilization. We selected fry at this early developmental stage before overt social interactions occurred (Fraley & Fernald, 1982) to ensure that the scope of our experiment captured meaningful social experience as early as possible during development. Individual broods were then placed in shallow water in a petri dish, and a digital image was taken with a ruler for scale. ImageJ was used to measure standard length (SL, mm) (Schneider et al., 2012), from the tip of the jaw to the caudal peduncle. Broods ranged in size from 4-21 fry (mean: 12.43 ± 2.84 fry, median: 15), and fry size ranged from 4.59-8.13 mm SL (mean: 5.96 ± 0.095 mm, median: 5.81 mm). See Supplemental Table 1 for mean, median, minimum, and maximum SL of fry per brood. All 7 broods were then placed briefly (∼30 min) in a common bucket to intermix. Given their developmental stage, social interactions were highly unlikely to occur, but visual, chemosensory, and tactile social sensory cues were likely exchanged. We used a hand net to remove individuals from the bucket and haphazardly assigned them to different home tanks and treatment groups (stable groups, socialized pairs, or stable pairs; see below for descriptions). Methylene blue was added to the water to reduce the incidence of fungal infection during this time. After 7 days, we confirmed visually that the fry were all mobile and their yolks had been fully absorbed. The sex ratio of fish in this experiment is not known because sex cannot be determined anatomically until reproductive maturation. The sex ratio of *A. burtoni* broods (a laboratory population descended from wild-caught fish) is approximately 1:1 (Heule et al., 2014).

### Early-life social experiences

Fry were reared for 32 or 35 days (depending on the behavioural test day, see below) in one of three conditions to manipulate early-life social experience: stable groups of 6 fish (n=6 groups), stable pairs of 2 fish (n=7 pairs), and socialized pairs of 2 fish (n=8 pairs) (Figure 1). These treatment groups were designed to manipulate the number of early-life social partners. In the stable groups, every individual had 5 social partners. In the stable pairs, every individual had 1 social partner. In the socialized pairs, 1 individual was removed from the tank using a hand net every 5 days and was replaced with a novel partner. The fish that was removed was transferred to another socialized pair tank with a novel partner. The social exchanges began on experimental day 12 (all fry were sufficiently mature on day 7, followed by 5 days with the first social partner). After a total of 4 exchanges, each of the fish in the socialized pairs were exposed to a total of 5 social partners, the same number of partners that each individual experienced in the stable groups. However, the size of the social group for socialized pairs was always 2 fish, the same as in the stable pairs. As a control, on the day that fish in the socialized pairs were transferred, we used a hand net to remove one fish from each of the stable pairs and stable groups. That fish was immediately returned to the same home tank so that group membership in the stable pairs and stable groups was consistent.

Stable and socialized pairs were housed in small acrylic tanks (23 × 15 × 15 cm), while the stable groups were housed in larger acrylic tanks (30 x 20 x 17 cm). One small terracotta pot (5 cm at the widest diameter) was placed in each tank, and an air stone was used to keep the water oxygenated. There was no substrate on the bottom of the tank. White plastic dividers were placed outside of the tanks to prevent visual contact between tanks. The fish were maintained on a 12:12 light:dark schedule at 24°C, and they were fed Hikari Middle Larval Stage Plankton (Pentair Aquatic Eco-Systems, Cary, NC) once a day. We mixed 0.015 g of fish food in 200 mL of water. The mixture was actively stirred, and a transfer pipette was used to add 1 mL to the pairs and 3 mL to the groups.

### Tagging individual fish

To track individuals as they developed over the course of the experiment, we tagged every fish following the first rotation in the socialized pairs. We waited until this time so the fish would be slightly bigger, as their very small size is a challenge for existing tagging methods (Lotrich & Meredith, 1974; Solomon-Lane & Hofmann, 2018; Thresher & Gronell, 1978). We first lightly anesthetized each fry in 0.03 g MS-222 / L aquarium water (buffered with sodium bicarbonate to pH 7.0 –7.5). They were then removed from the water and placed on a wet paper towel. We placed a small amount of Alcian Blue (Fisher Scientific, Pittsburgh, PA) powder onto the dorsal muscle of the fish and perforated the surface of the fish using a pulled capillary tube micropipette. This method of tattooing allowed the ink to seep into the tissue of the fish. We used different tag locations (left or right; anterior, middle, or posterior regions) on the dorsal muscle to differentiate individuals in the same tank. In the socialized pairs, tagging was sufficient in most tanks to identify and move the same individual each time. The persistence of the tag varied, and in the best cases, it remained visible for 3 weeks. If the tags were not visible in the socialized pair, the smaller individual was moved.

### Survival and number of early-life social partners

The tagging procedure had an 85% survival rate. Fish that did not survive tagging or that died at another point in the experiment were replaced with a fish of similar size and age, selected from a community tank. The replacement fish were not included in the individual behaviour assays conducted at the end of the experimental rearing period (see below). We tracked the exact number of social partners for each experimental tank. All fish in the socialized pairs had 5 social partners. Stable group fish had an average of 6.43 ± 0.14 social partners. Stable pair fish had an average of 1.2 ± 0.13 social partners. In our analyses, we consider the treatment groups as categorical, although there is limited variation in the number of social partners (Supplemental Figure 1).

### Behavioural Observations

<u>Home tanks</u>: We recorded social behaviour in the home tanks twice during the rearing period, on days 20 and 25 of the experiment, at the same time of day. Cameras (Warrior 4.0, Security Camera Warehouse, Asheville, NC) were positioned above the tanks, and with the tank lids removed, all areas of the tank were visible except under the terracotta pot. Using Solomon Coder (www.solomoncoder.com), 10 minutes of video was scored for approaches and displacements. Approaches were counted anytime one fish swam directly towards any part of another fish, within 3 body lengths. Approaches can range from affiliative to aggressive, and the response of the approached fish provides insight into the degree of agonism. If the approached fish swam away in any direction, it was scored as a displacement, which is an indication that the approach was aggressive. Because the identification tags were not visible on the video, we counted the total number of social interactions (approaches, displacements) for the group or pair. We also quantified the number of times a fish entered the terracotta pot territory, defined as at least 50% of a fish’s body crossing into / under the pot. Exiting the territory was highly correlated with entering the territory (p<0.0001, r^2^=0.97); therefore, we only present data for entering. To measure social grouping, we captured screenshots from the video every 30 s for a total of 20 time points. Distance was measured between every dyad, from a focal point in the centre of the head, using ImageJ (Schneider et al., 2012). The researchers scoring these behaviours were unaware of the treatment of the pairs (socialized vs. stable), but the stable groups were easily recognizable. For behaviour and distance, we analysed the raw values (behaviours per minute and distance), as well as scaled these values to account for differences in group size (behaviours per minute divided by 2 for the pairs and divided by 6 for the groups) and tank size (distance divided by the hypotenuse of the tank) among treatments.

<u>Individual Behaviour Assays</u>: After the 32 or 35-day rearing period, we measured individual behaviour in a series of assays, including open field exploration, social cue investigation, dominant behaviour, and subordinate behaviour. Every fish in the socialized (final sample size: n=9) and stable pairs (final sample size: n=10) was included in the individual assays. Three of the 6 fish from each stable group were selected randomly to be included in the individual assays using a random number generator (final sample size: n=14). The procedures have been described previously in detail (Solomon-Lane & Hofmann, 2019). Briefly, these assays (or similar) are used across species to assess different elements of behavioural phenotype, including behaviours affected by early-life social experience (Sih, 2011; Sih et al., 2015; Solomon-Lane & Hofmann, 2019; Taborsky, 2016). The open field exploration is used to assess locomotion and anxiety-like behaviours (e.g., Cachat et al., 2010; Prut & Belzung, 2003). The social cue investigation is used as a measure of social motivation or preference (e.g., Bonuti & Morato, 2018; Moy et al., 2004). The dominant and subordinate behaviour assays are used to assess how the focal fish behaves when in the position to be dominant or subordinate. In social groups, including for juvenile *A. burtoni* (Solomon-Lane et al., 2022), most individuals will be subordinate to some group members and dominant over others at a given time (e.g., Chase et al., 2022). Social status also changes over a lifetime (Fernald & Maruska, 2012; Hofmann, 2003).

The behavioural assays took place in novel experimental aquaria, which were the same size as the pairs’ home tanks (3.27 L, 23 × 15 × 15 cm). Cameras recorded behaviour from above, and all areas of the tank were visible except under the terracotta pot. A permanent marker was used to delineate different zones (Figure 1B): a territory zone that contained a small terracotta pot, a close zone, a far zone, and an investigate zone where the social cue was placed for the social cue investigation assay. Each assay lasted for 30 min. The focal fish was alone in the tank for the open field exploration (minutes 20-30 were analysed). For the social cue investigation, a live, novel cue fish inside of a 20 mL glass vial was placed in the investigate zone (see placement in Figure 1B, minutes 2-12 were analysed). In the open field and social cue assays, we recorded the number of times a fish entered a zone and how long it spent there. After removing the vial and cue fish, a free-swimming, novel fish that was smaller than the focal fish was added to the tank to assess dominant behaviour (minutes 2-12 were analysed). After removing the small fish, a free-swimming novel fish larger than the focal was added to assess subordinate behaviour (minutes 2-12 were analysed). In the dominant and subordinate behaviour assays, we scored approaches and displacements. The researchers scoring these behaviours were unaware of the treatment of the focal fish.

From the approaches and displacements in the dominant and subordinate behaviour assays, we calculated two additional social measures using the *compete* package in R (Curley, 2016). First, we calculated an adjusted version of David’s Score, which is a dominance index. Rather than categorizing all social interactions as a one-dimensional “win” or “loss,” we incorporated both approaches and displacements, such that the agonistic outcome of individual i in interaction with another individual j is the number of times that i displaces j, divided by the total number of interactions (approaches) between i and j (i.e., i displaces j / (i approaches j + j approaches i)). The rest of the calculations were done as described in Gammell et al. (2003). See the Supplemental Information in Solomon-Lane et al. (2022) for the R code. Second, we calculated directional consistency index (DCI) to assess whether patterns of approaching in the dyads are reciprocal, from perfectly reciprocal (0) to unidirectional (1). We also used the *compete* package to run a randomization test to determine if the directionality was significantly more asymmetrical (unidirectional) than expected by chance (Leiva et al., 2008).

### Water-borne cortisol

Following the individual behaviour assays, we collected water-borne hormone samples. This is a non-invasive method of quantifying steroid hormones, and values strongly correlate with circulating hormone levels (Kidd et al., 2010). We chose cortisol, as HPI axis function is often affected by early-life experience (e.g., Champagne & Curley, 2005; Taborsky et al., 2013) and cortisol can correlate with behaviour and behavioural syndromes (e.g., Koolhaas et al., 1999). Focal fish were placed individually in clean glass vials filled with 15 mL of clean aquarium water for 2 hrs. Visual barriers were placed around the vials to limit disturbances to the fish during the collection. After two hours, fish were removed from the vial using a hand net rinsed in clean aquarium water, and samples were frozen at 4°C until processing. Hormones were extracted from the water samples using 3 cc Sep-Pak Vac C18 columns (Water Associates, Milford, MA). Columns were first primed by passing 2 mL methanol through the columns twice, being careful not to let the columns run dry. This was followed by running 2 mL ultrapure water through the columns twice. The thawed samples were then passed through the columns from the sample vials using the vacuum pump. Ultrapure water (2 mL twice) was then passed through the columns again, followed by 5 min of the vacuum to dry the columns. Finally, the hormone samples were eluted into 13 x 100 borosilicate test tubes by passing 2 mL methanol through the columns twice. The vacuum was used again to dry the columns. The sample in methanol was then evaporated under a gentle stream of nitrogen at 37°C and resuspended to a volume of 200 μL per sample (5% EtOH and 95% ELISA buffer). Resuspended samples were shaken on a multitube vortex for 1 hr before being stored at -20°C. Before measuring cortisol, samples were thawed and shaken again on the multitube vortex for 1 hr. ELISAs were completed according to the supplier’s instructions (Cayman Chemical, Ann Arbor, Michigan).

Cortisol data are presented here as pg hormone per mL sample volume per hr sample collection per g body weight. During hormone collection, fish excrete hormones into the water through their gills, urine, and faeces. The gills of larger fish have a larger surface area compared to smaller fish, thus body size can influence water-borne hormone levels. Fish size was measured following the collection of water-borne hormones. Mass was measured on an analytical balance, and SL was measured using ImageJ (Schneider et al., 2012) from digital images with a rule for scale. We calculated the condition index as Fulton’s *K*, which estimates expected mass as SL^3^. We used the mass and SL of 400 laboratory *A. burtoni* across a range of ages and sizes to empirically calculate expected mass as: mass=0.00002*SL^3.02^ (Stevenson & Woods, 2006).

### Ethical Note

All research was done in compliance with the Institutional Animal Care and use Committee (IACUC Protocol # 19-001). We used non-invasive approaches when possible, such as collecting water-borne hormones, and we used a tested and effective anaesthetic (MS-222) for tagging to minimize stress and pain. We took steps to minimize stress throughout the experiment, including limiting handling time to less than 2 min when measuring size or transferring between tanks. A total of 76 juvenile *A. burtoni* were included in this study, and fish were returned to community tanks at the end of the experiment.

### Statistical Analyses

All statistical analyses were conducted using R Studio (R version 4.2.1) (RStudio Team, 2022). Results were considered significant at the p<0.05 level, and averages ± standard error of the mean are included in the text. The boxes of the box and whisker plots show the median and the first and third quartiles. The whiskers extend to the largest and smallest observations within or equal to 1.5 times the interquartile range. To test for the effects of early-life social experience, we compared body size and condition index; behaviour in the home tank rearing environments (total behaviours per minute and behaviours per fish per minute); spatial distancing in the home tank rearing environments (raw distance and scaled to account for the different sized aquaria for pairs vs. groups); behaviour in the open field exploration and social cue investigation (frequency entering and time in the territory, close, far, and investigate zones); social behaviour (approaches, displacements, approaches received, submissions) and status (David’s score, DCI) in the dominant and subordinate assays; cortisol; and principal components (see below, PCs 1-5) across fish reared in the stable groups, socialized pairs, or stable pairs. For data that met the assumptions of parametric statistics, we used one-way ANOVAs. The cortisol data were log transformed first to meet the assumption of normal distribution. *Post hoc* analysis of significant ANOVA results was calculated using Tukey HSD tests. For data that did not meet the assumptions of parametric statistics, we used non-parametric Kruskal-Wallis tests, and Dunn’s tests with Bonferroni correction were used for *post hoc* analysis of significant results. Eta squared is reported for the effect size of both the one-way ANVOAs and Kruskal-Wallis tests (small effect: 0 < η2 < 0.01; moderate: 0.01 < η2 < 0.06; large: 0.06 < η2). We used Principal Components Analysis (PCA) to identify how behaviour in the open field exploration, social cue investigation, dominance behaviour assay, subordinate behaviour assay, and cortisol clustered.

## RESULTS

### Body size and condition

To determine whether early-life experience in a stable group, socialized pair, or stable pair affected body size and condition, we compared SL, mass, and condition index across treatment groups. We found that mass differed significantly across treatment groups (χ^2^_2_=6.39, p=0.041, *η*^2^=0.15; Supplemental Figure 2A). *Post hoc* analysis showed juveniles reared in stable pairs were significantly heavier than those from stable groups (p=0.036). There were no significant differences in mass between stable groups and socialized pairs (p=0.63) or between stable pairs and socialized pairs (p=0.82). There were no significant treatment differences in SL (F_2,30_=2.15, p=0.14; Supplemental Figure 2B). Because we found an effect of rearing experience on mass, but not SL, we also asked whether condition index was affected by treatment. We found no significant treatment differences in condition index (F_2,30_=0.45, p=0.62, Supplemental Figure 2C). Mass was also positively correlated with water-borne cortisol levels (p=0.017, r^2^=0.15, Supplemental Figure 2D); therefore, we corrected for body size in the cortisol analyses below.

### Home tank social behaviour

We observed juveniles in the stable groups, socialized pairs, and stable pairs to understand the differences in social experience and environment among the rearing treatments. Unsurprisingly, there were significantly more approaches and displacements in groups compared to pairs (both stable and socialized), and there were significantly more entrances to the territory in groups compared to stable pairs (Figure 2A). There were no differences in efficiency (Figure 2B). There were also no differences in rates of behavioural interactions per fish. Mean dyad distance was significantly larger in groups than pairs (Figure 2C), but interestingly, the minimum dyad distance was significantly smaller in groups than socialized pairs (Figure 2D). There were no treatment differences in mean dyad distance when scaled for tank size. Statistics are reported in Table 1 and Supplemental Table 2.

**Figure 2:**
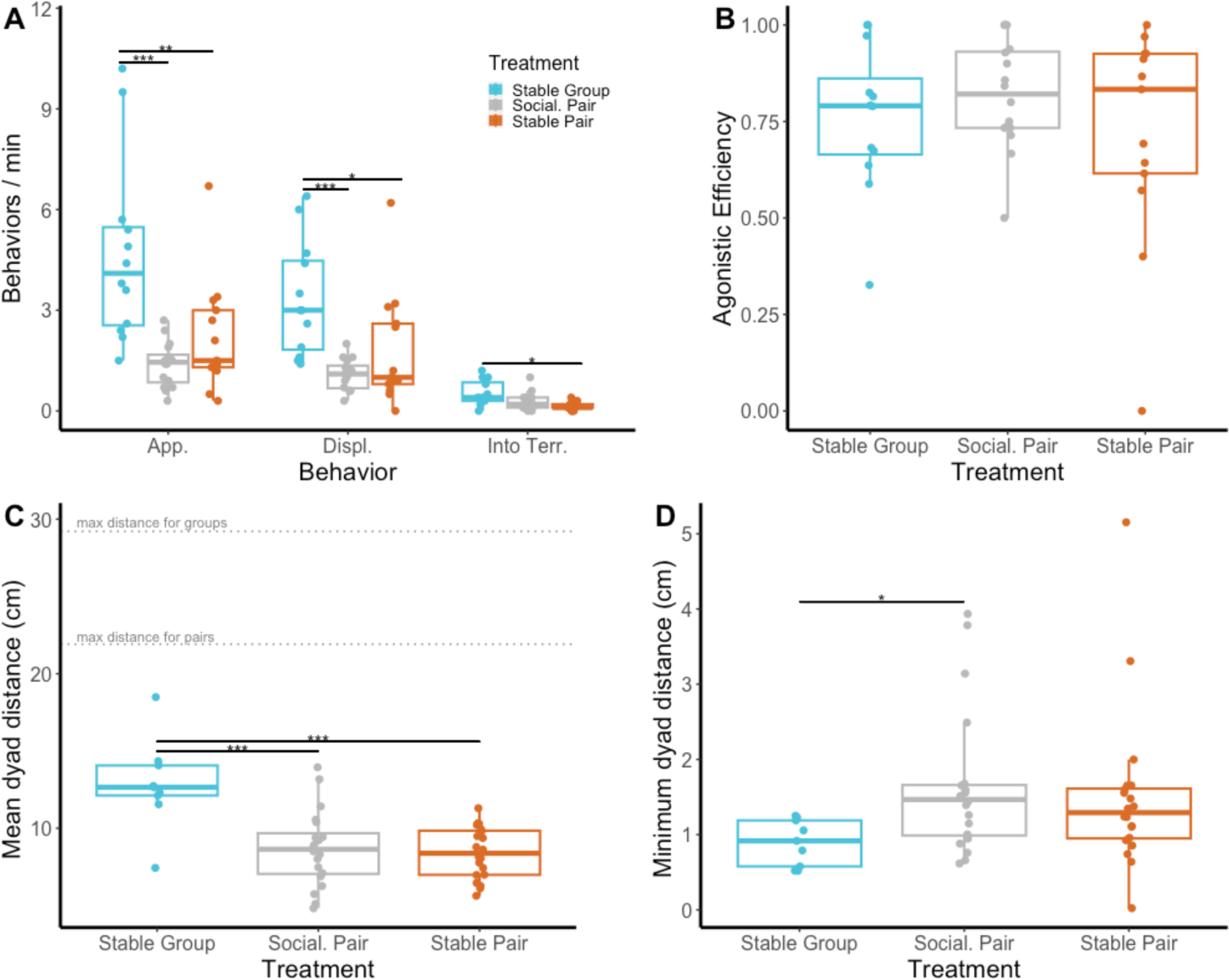
Social behaviour and distances in the stable groups, socialized pairs, and stable pairs. A) Frequency of social approaches, displacements, and times entering the territory / terracotta pot in the rearing environments. B) Total agonistic efficiency (total displacements / total approaches). C) Distance between all dyads of fish was measured twice for each tank (20 time points each, 40 total time points). Mean distance between dyads was then calculated per tank (each data point is the mean for one tank). The maximum distance lines are the lengths of the diagonal along the bottom of the tanks. D) The minimum distance recorded between a dyad across all time points. *p < 0.05, **p < 0.01, ***p<0.001.

**Table 1:** Treatment differences in social behaviour and distances between dyads in the stable group, socialized pair, and stable pair home tanks.

| Behaviour / measure | DF | Test Statistic | p-value | Effect size | *Post hoc* / direction of effect |
| --- | --- | --- | --- | --- | --- |
| Approaches | 2, 38 | F=11.46 | 0.00013 | 0.38 | **Stable group > social. pair: p<0.0001**<br>**Stable group > stable pair: p=0.0061**<br>Social. pair vs. stable pair: p=0.41 |
| Displacements | 2, 37 | F=11.37 | 0.00014 | 0.38 | **Stable group > social. pair: p<0.0001**<br>**Stable group > stable pair: p=0.02**<br>Social. pair vs. stable pair: p=0.21 |
| Efficiency | 2 | $\chi^2=1.06$ | 0.59 | | |
| Into territory | 2 | $\chi^2=10.81$ | 0.0045 | 0.23 | **Stable group > stable pair: p=0.0037**<br>Stable group > social. pair: p=0.071<br>Social. pair vs. stable pair: p=0.75 |
| Mean dyad distance | 2, 46 | F=13.62 | <0.0001 | 0.37 | **Stable group > stable pair: p<0.0001**<br>**Stable group > social. pair: p<0.0001**<br>Social. pair vs. stable pair: p=0.92 |
| Minimum dyad distance | 2 | $\chi^2=7.49$ | 0.0025 | 0.12 | **Stable group > social. pair: p=0.021**<br>Stable group > stable pair: p=0.085<br>Social. pair vs. stable pair: p=1.0 |

Results of one-way ANOVAs (F) or nonparametric Kruskal-Wallis tests (χ^2^). Eta-squared is reported for effect sizes. Significant results in bold. Tukey HSD tests were used for *post hoc* analysis of significant ANOVA results. Dunn’s tests were used for *post hoc* analysis of significant Kruskal-Wallis results.

### Open field exploration and social cue investigation

For the open field exploration and social cue investigation, we compared across treatment groups for the frequency of entering each zone of the tank, as well as time spent in each zone. Statistics are reported in Table 2 (Supplemental Figure 3). Overall, we found for the open field exploration that fish from the socialized pairs entered the territory zone significantly more frequently than stable pair fish, and fish from the stable groups spent significantly more time in the far zone than stable pair fish. In the social cue investigation, we found that social group fish entered the far zone significantly more frequently and spent significantly more time in the territory and far zones, than fish reared in stable pairs.

**Table 2:**
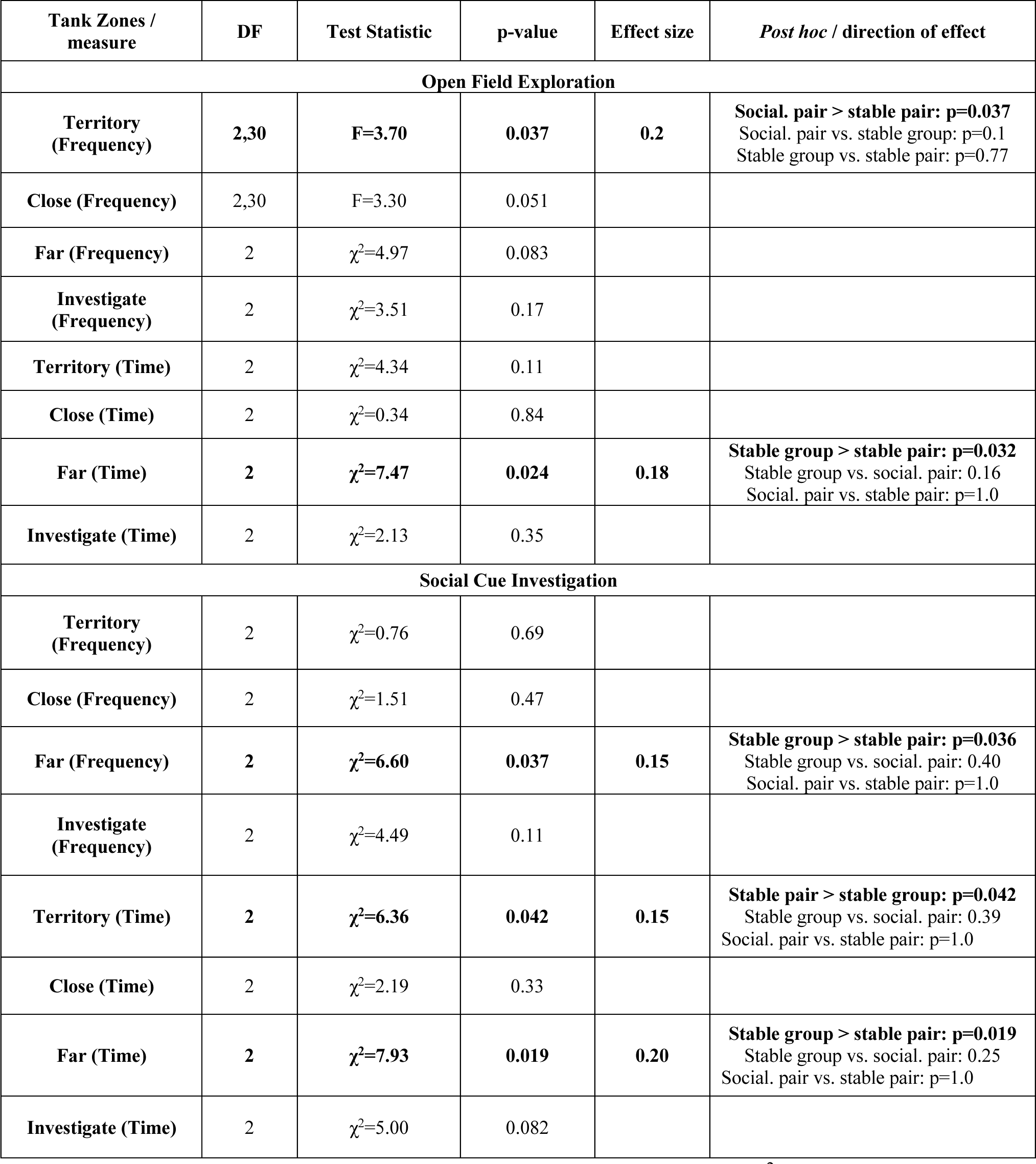
Treatment differences in the frequency entering and time spent in tank zones during the open field exploration and social cue investigation.

Results of one-way ANOVAs (F) or nonparametric Kruskal-Wallis tests (χ^2^). Eta-squared is reported for effect sizes. Significant results in bold. Tukey HSD tests were used for *post hoc* analysis of significant ANOVA results. Dunn’s tests were used for *post hoc* analysis of significant Kruskal-Wallis results.

### Dominance and Subordinate Assays

To determine whether rearing experience affected social behaviour and status, we quantified patterns of social interaction between the focal fish and a novel social partner (cue fish). Because relative physical size often affects dominant and subordinate social dynamics for juveniles (Solomon-Lane et al., 2022), pairing the focal fish with a smaller fish (dominance assay) presented the focal fish with the opportunity to express social behaviours as the dominant. In pairing the focal fish with a larger fish (subordinate assay), the focal fish can express social behaviours as the subordinate. We found significant differences in social behaviour and status in both the dominance and subordinate assays (statistics reported in Table 3, Figure 3). In the dominance assay, socialized pair fish approached and displaced significantly more than stable pair fish, and their David’s Scores were significantly higher than both stable pair and stable group fish. In the subordinate assay, socialized pair fish received significantly more approaches than stable pair fish from the larger cue fish with which they were paired. Socialized pairs with their cue fish also had significantly lower directional consistency than stable group or stable pair fish.

**Figure 3:**
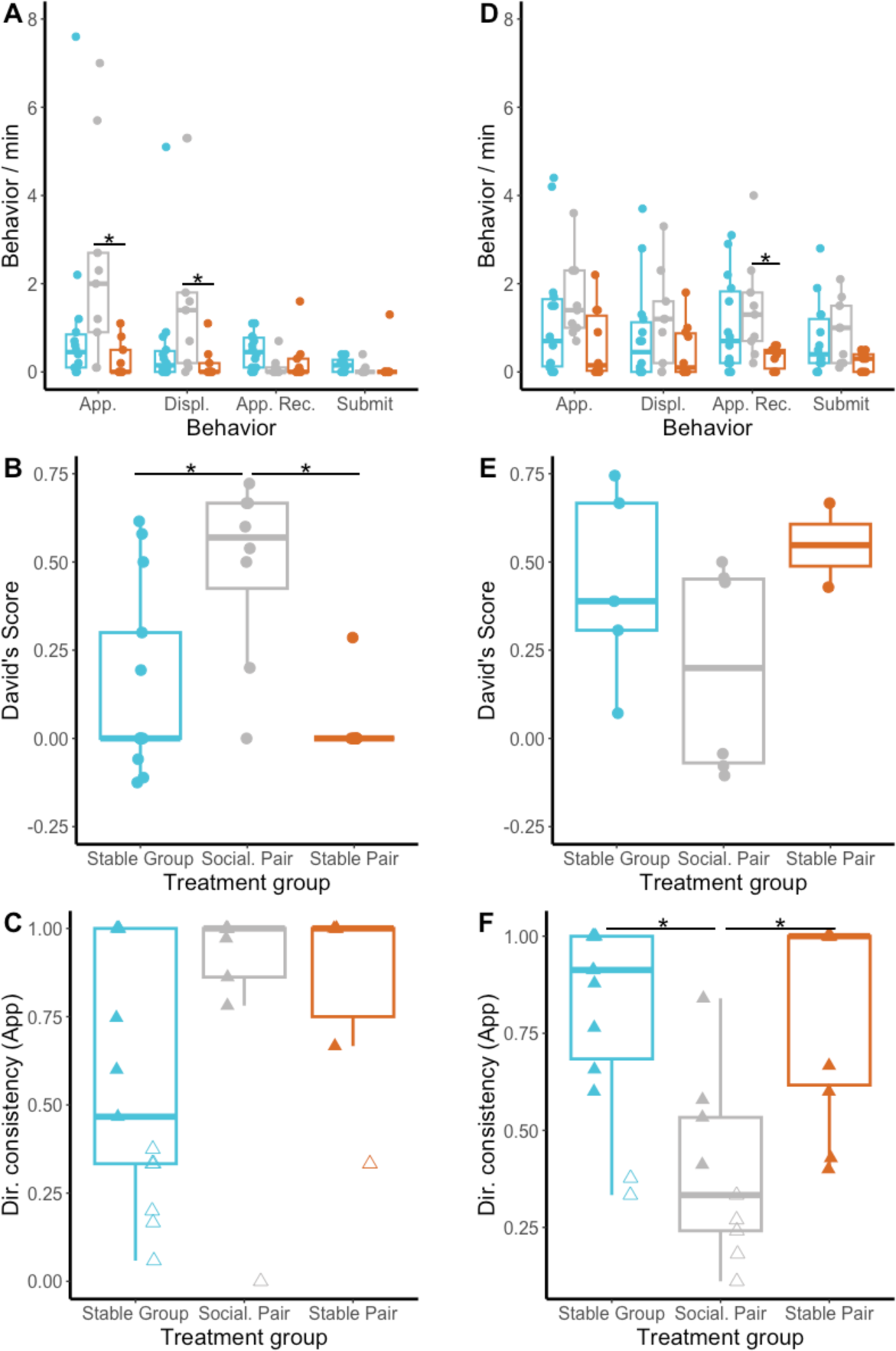
Social measures from the dominant (A-C) and subordinate (D-F) behaviour assays. A) Social approaches (App.), displacements (Displ.), approaches received (App. Rec.), and submissions (Submit) in the dominance behaviour assay. B) Focal fish David’s score in the dominance behaviour assay. C) Directional consistency (calculated based on approaches) in the dominance behaviour assay. Filled triangles indicate directionality was significantly greater than zero. Empty triangles indicate pairs were not significantly directional. D) Social behaviour in the subordinate behaviour assay. B) Focal fish David’s score in the subordinate behaviour assay. C) Directional consistency (calculated based on approaches) in the subordinate behaviour assay. *p<0.05.

**Table 3:**
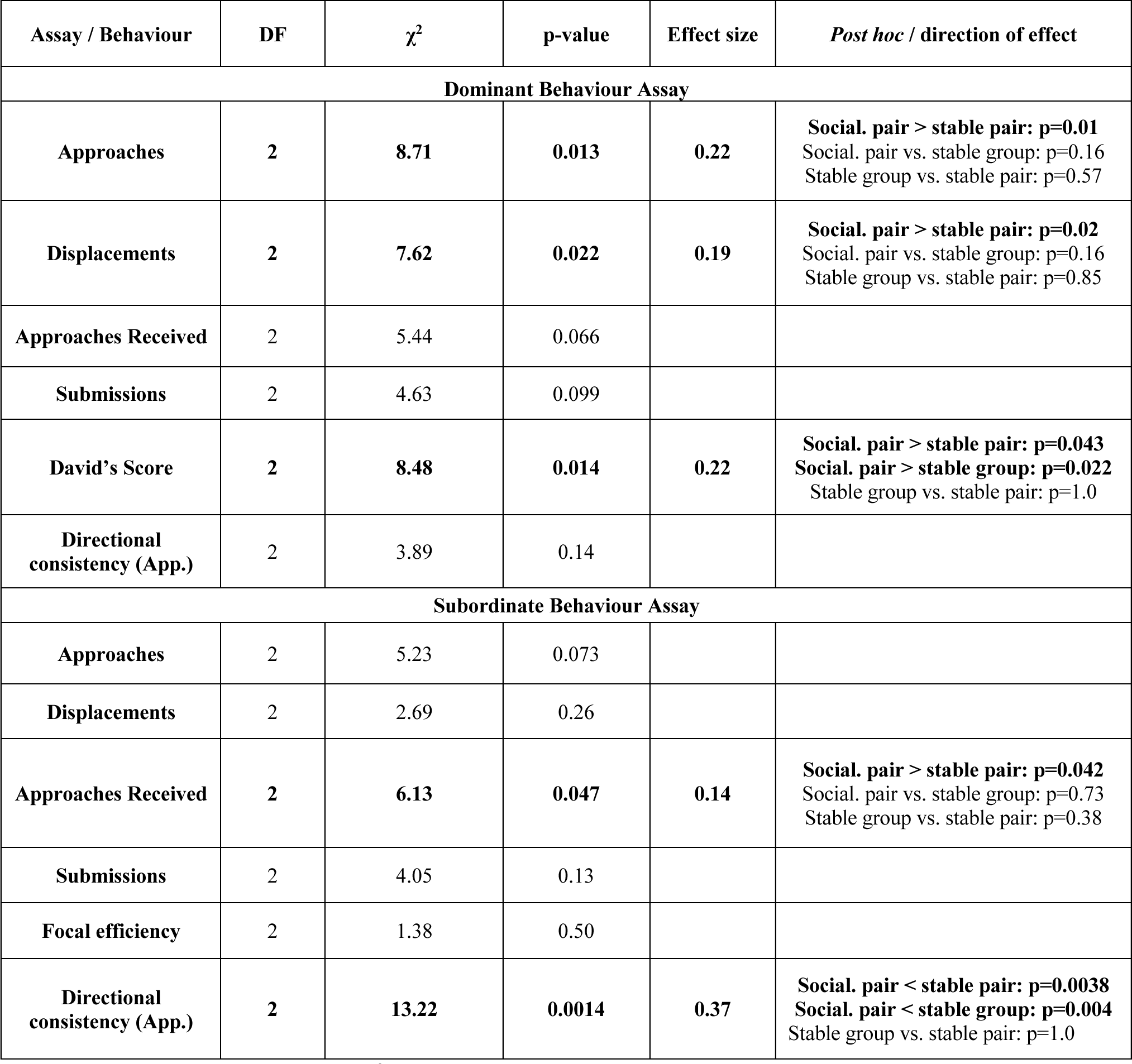
Treatment differences in social behaviour and status in the dominant and subordinate behaviour assays.

Results of Kruskal-Wallis tests (χ^2^). Eta-squared is reported for effect sizes. Significant results in bold. Dunn’s tests were used for *post hoc* analysis of significant results.

### Cortisol

Following the behaviour assays, we collected water-borne cortisol to determine whether hormone levels were affected by rearing experience and/or associated with behavioural phenotype (see PCA below). We found there were no significant differences in cortisol across treatments (F_2,29_=2.30, p=0.12, Figure 4).

**Figure 4:**
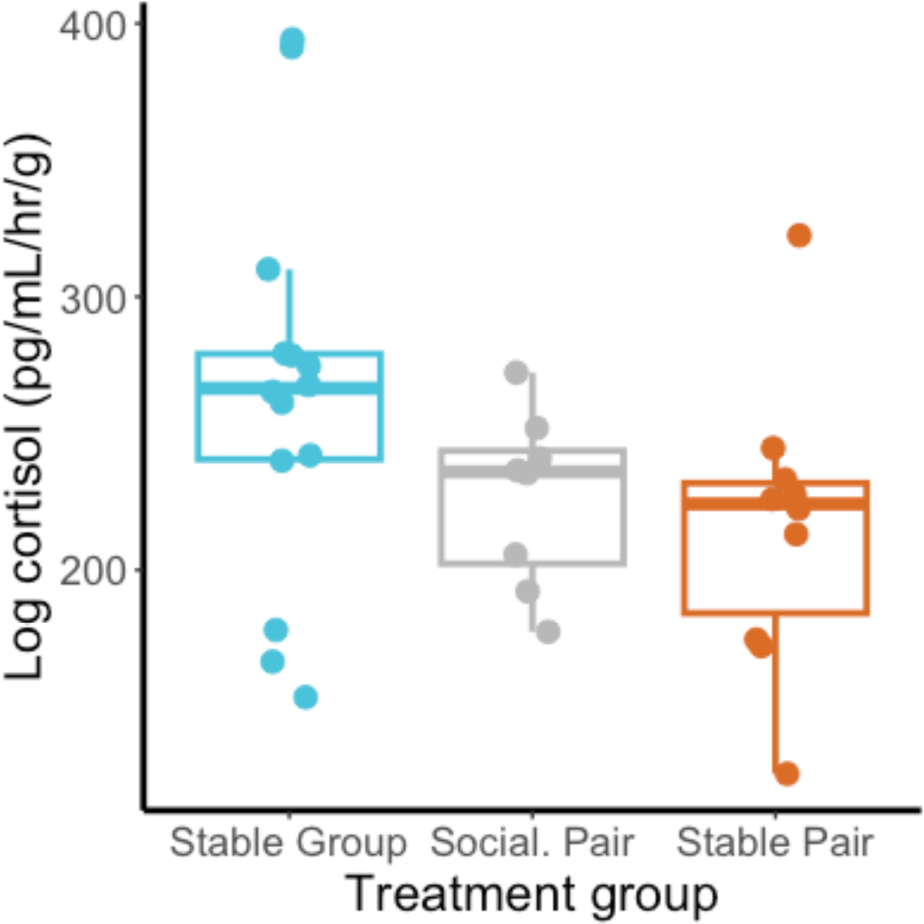
Focal fish water-borne cortisol (pg/mL/hr), corrected for body mass (g). Log transformed data are shown.

### Integrative analysis of behaviour, social status, and cortisol

Given the treatment differences we identified across the open field exploration, social cue investigation, dominance assay, and subordinate assay, we next used PCA to integrate across assays. This allowed us to identify subsets of factors that together contribute to juvenile phenotype and test if suites of factors—principal components (PCs)—differed significantly across treatment groups. To determine which variables to include from the open field exploration and social cue investigation (time and frequency in the territory, close, far, and investigate zones), we ran an initial PCA with just these variables. Examining the vector plot for PC1 with PC2 (Supplemental Figure 4), we found that nearly all of the open field exploration measures were strongly aligned with the same measure in the social cue investigation (i.e., the vectors for open field and social cue time in the territory zone are identical). The exceptions were for time spent in the far and investigate zones. Therefore, to avoid unnecessary replication, we chose to include all of the variables from the social cue investigation, in addition to time spent in the far and investigate zones during the open field exploration. From the dominant and subordinate assays, we included approaches, displacements, approaches received, submissions, and David’s score. Finally, we included cortisol.

Here, we focused on the first 5 PCs, which together explain 76.6% of the variation in the data (Figure 5A). We found significant effects of rearing experience on PC1 (25.1%) and PC5 (8.48%). For PC1, stable pairs were significantly higher than stable groups, and there was a trend for stable pairs to be higher than socialized pairs (p=0.063) (Figure 5B). Behaviours from the dominant behaviour assay and the social cue investigation assay loaded strongly on PC1. Approaches and displacements when in a dominant social role; time in and frequency entering the far and investigate zones during the social cue investigation; and frequency entering the close zone during the social cue investigation loaded strongly in the same direction. Time spent in the territory zone during the social cue investigation loaded strongly in the opposite direction. For PC5, socialized pairs were significantly higher than both stable groups and stable pairs (Figure 5C). David’s Score from the dominance assay and time in the investigate zone during the social cue investigation loaded strongly together in the same direction. In the opposite direction, submissions as a dominant; cortisol; time and frequency in the close zone during the social cue investigation; and the frequency entering the territory zone during the social cue investigation loaded strongly together. Statistics for the PC1-PC5 treatment comparisons are in Table 4 (also see Supplemental Figure 5).

**Figure 5:**
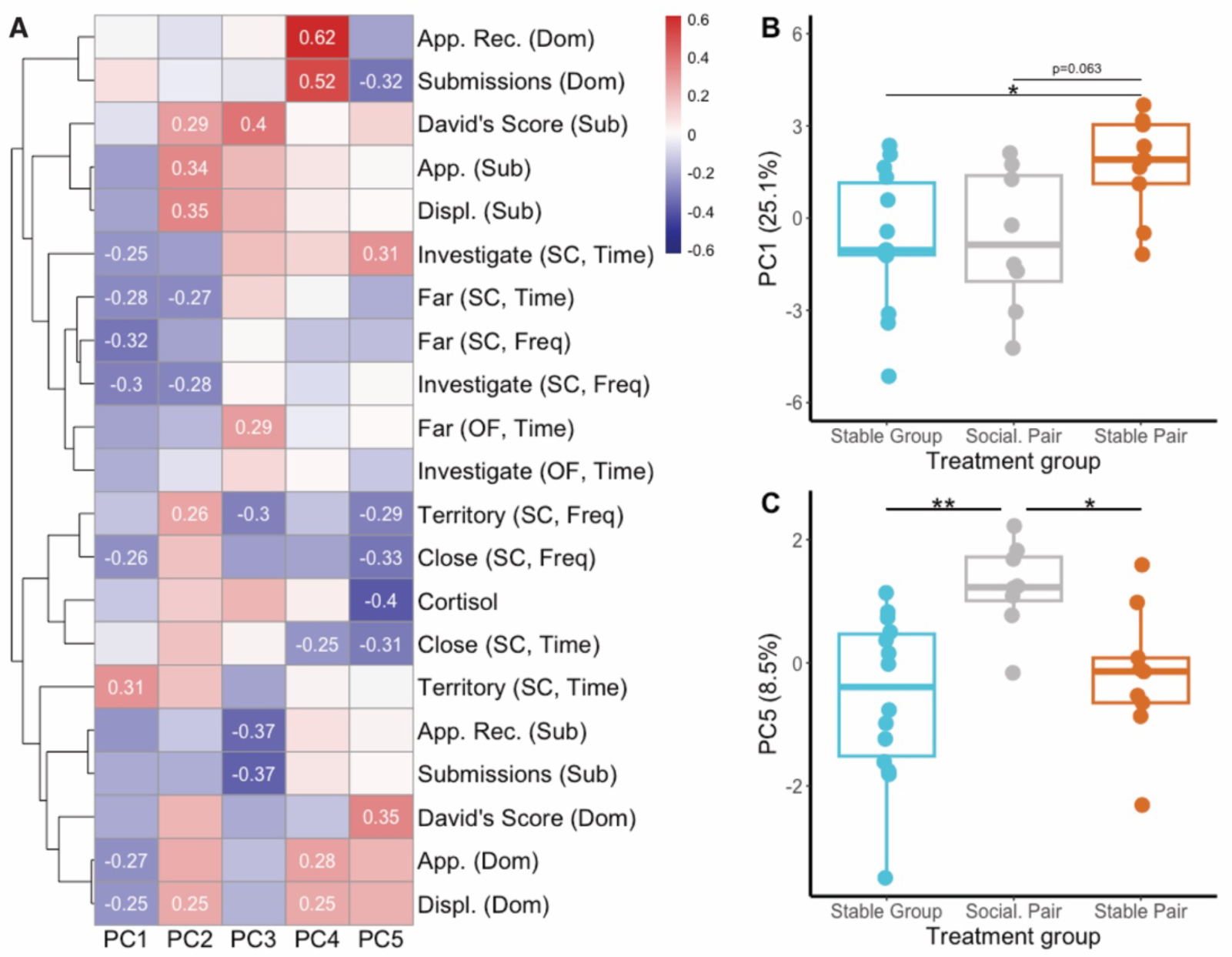
Principal components analysis (PCA) of cortisol and behaviour—including open field (OF) time in the far and investigate zones; social cue (SC) time in and frequency entering the territory, close, far, and investigate zones; and dominant and subordinate behaviour assay approaches (App.), displacements (Displ.), approaches received (App. Rec.), submissions, and David’s Score. A) A heatmap of eigenvalues showing the PCA variables that load on PC1 (25.1%), PC2 (18.2%), PC3 (14.3%), PC4 (10.5%), and PC5 (8.5%). Numerical values are shown for variables stronger than ±0.25. Rows are hierarchically clustered. B) Treatment differences in PC1. C) Treatment differences in PC5. Percentages refer to the amount of variance explained by that PC. *p<0.05. **p<0.01.

**Table 4:** Treatment differences for principal components (PC) 1-5.

| Principal component<br>(% variation) | DF | Test Statistic | p-value | Effect size | *Post hoc* / direction of effect |
| --- | --- | --- | --- | --- | --- |
| **PC1 (25.1%)** | 2,28 | **F=4.19** | **0.026** | **0.23** | **Stable pair > stable group: p=0.032**<br>Stable pair > social. pair: p=0.063<br>Stable group vs. social. pair: p=1.0 |
| **PC2 (18.2%)** | 2,28 | F=1.82 | 0.18 |  |  |
| **PC3 (14.3%)** | 2,28 | F=0.97 | 0.39 |  |  |
| **PC4 (10.5%)** | 2 | $\chi^2=2.69$ | 0.26 | | |
| **PC5 (8.5%)** | 2,28 | **F=6.62** | **0.0044** | **0.32** | **Social. pair > stable group: p=0.0036**<br>**Social. pair > stable pair: p=0.036**<br>Stable group vs. stable pair: p=0.75 |

Results of one-way ANOVAs (F) or nonparametric Kruskal-Wallis tests (χ^2^) for PC1-PC5.

Percentages indicate the variance explained. Eta-squared is reported for effect sizes. Significant results in bold. Tukey HSD tests were used for *post hoc* analysis of significant ANOVA results. Dunn’s tests were used for *post hoc* analysis of significant Kruskal-Wallis results.

## DISCUSSION

We investigated the effects of early-life social experience, in the first month of life, on social behaviour and neuroendocrine stress axis function in juvenile *A. burtoni*. We tested the hypothesis that the number of novel social partners experienced during early-life is a behavioural mechanism driving variation in development. In manipulating the number of social partners, our experimental treatment groups—stable groups, socialized pairs, and stable pairs—also varied in social group size (pairs vs. group) and group stability (socialized vs. stable). We present strong evidence for early-life social effects on juvenile social behaviour, consistent with previous studies in this species (Fernald & Hirata, 1979; Fraley & Fernald, 1982; Solomon-Lane & Hofmann, 2019). In particular, our results support two behavioural mechanisms: the number of social partners and social stability. Although early-life social effects are widely documented across social species, the specific attributes of social experience that affect the mechanisms and trajectory of developmental plasticity are not typically identified (Kasumovic, 2013; Taborsky, 2016). The social interactions and social spacing we observed in the home tanks also provide insights into how social environments vary within and across treatments and may shape the experiences individuals accrue during development. Causation at this level is needed because it is these environmental elements that interact with genes dynamically over development to influence plasticity and the emergence of adult phenotype (Kasumovic, 2013; Taborsky, 2016, 2017). Overall, it is likely that multiple behavioural mechanisms contribute and interact to shape development (e.g., Branchi et al., 2013) and adult phenotype.

We found that manipulating early-life social experience affected behaviour across multiple contexts, including the open field exploration, social cue investigation, dominant behaviour assay, and subordinate behaviour assay. Principal components analysis revealed that these effects were correlated across contexts. Principal component 1 (25.1%) and PC5 (8.5%), which both differed significantly across treatment groups, had behaviours from multiple assays that loaded strongly. These results support our previous finding that juvenile *A. burtoni* behaviour can form a syndrome, and an individual’s position along the syndrome continuum is sensitive to early-life social experience (Solomon-Lane & Hofmann, 2019). A syndrome is a population-level measure in which rank-order differences between individuals are correlated across contexts and/or over time (Bell, 2007). Behavioural syndromes have been identified across species and can indicate consistency in individual behaviour across contexts and/or over time (Bell, 2007; Sih, Bell, & Johnson, 2004; Sih, Bell, Johnson, et al., 2004). The PC1 syndrome included approaches and displacements in the dominance assay loading strongly in the same direction with time in and frequency entering the investigate and far zones, and frequency entering the close zone, during the social cue investigation. Time in the territory zone during the social cue investigation loaded in the opposite direction. This syndrome is highly similar to the one we identified after rearing juvenile *A. burtoni* in pairs or social groups of 16 fish (Solomon-Lane & Hofmann, 2019).

The treatment differences in PC1 are most consistent with an effect of the number of social partners. Stable pairs had significantly higher PC1 values compared to stable groups, and there was a strong trend to be higher than socialized pairs (p=0.063). There were no differences between stable groups and socialized pairs. Juveniles that experienced more social partners during early-life were more active during the social cue investigation, spent less time in the territory in the presence of a social cue, and were more socially interactive in a dominant social role. The direction of this effect is also the same as in our previous study: group-reared juveniles were more active and socially interactive than pair-reared juveniles (Solomon-Lane & Hofmann, 2019). This suggests that social experiences resulting from more novel partners may be an important behavioural mechanism underlying the effect of group size. Syndromes involving activity and social interaction are common across species (e.g., Conrad et al., 2011; Näslund & Johnsson, 2016) and may also be related to bold-shy and proactive-reactive behaviours (Bell, 2007; Conrad et al., 2011; Groothuis & Carere, 2005; Koolhaas et al., 1999; Sih, Bell, Johnson, et al., 2004).

It is not yet known how variable the number of social partners is for juvenile *A. burtoni* in the wild or in larger, more naturalistic laboratory social groups. Evidence from a diversity of other species suggests there can be considerable individual variation. In many species, siblings (and parents) are the most proximate—and sometimes the only—early-life social contacts. The number of offspring produced can vary both among individuals and within individuals across reproductive bouts / over time. Within a group or population, individuals can also vary in the size and makeup of their social network or niche (Barale et al., 2015; Beirão-Campos et al., 2016; Branchi et al., 2013; Croft et al., 2005; Förster & Cords, 2005; Maguire et al., 2021; Monclús et al., 2012; Pike et al., 2008; Saltz et al., 2016; Weinstein et al., 2014; Weinstein & Capitanio, 2008, 2008). For juveniles, this can have long-term, phenotypic effects (Branchi et al., 2013; Monclús et al., 2012; Weinstein et al., 2014; Weinstein & Capitanio, 2008). Socialization strategies are also used by humans with animals, such as working dogs (Gfrerer et al., 2018), family dogs (Howell et al., 2015), and livestock like piglets (Morgan et al., 2014; Salazar et al., 2018). Although many studies have manipulated the number of early-life social partners as a consequence of group size, group size could exert independent or interacting effects on phenotype. In studies that also controlled for group size, brown-headed cowbirds (*Molothrus ater*) were housed in stable or dynamic flocks, in which flock members were exchanged multiple times with novel birds. Dynamic flock males had more variable social networks over time, larger signing networks, and outcompeted stable flock males in mating opportunities (White et al., 2010). When housing conditions were later reversed, the new dynamic flock males still had higher reproductive success, which was achieved via changes in social strategy (Gersick et al., 2012). For juvenile *A. burtoni* in pairs and triads, individuals are not equally likely to initiate interactions. Both group size and relative body size influenced social group structure (Solomon-Lane et al., 2022). This suggests that, like adults (Maguire et al., 2021), social network position and connectivity may vary considerably across individuals, with consequences for developmental plasticity and behavioural development.

The treatment differences in PC5 (8.5%) support an effect of early-life social stability. Socialized pairs had significantly higher PC5 values than stable groups and stable pairs, which were not different from each other. David’s score in the dominance behaviour assay loaded in the same direction as time in the investigate zone during the social cue investigation. Submissions in the dominance behaviour assay, frequency entering the territory zone in the social cue investigation, time in and frequency entering the close zone in the social cue investigation, and water-borne cortisol loaded together in the opposite direction. Socialized pairs were more dominant in the dominance behaviour assay, spent less time in and near the territory zone, spent more time in the investigate zone, and had lower cortisol levels than stable groups or pairs. This suite of behaviours for PC5 shares multiple similarities with the PC1 syndrome, suggesting these syndromes may not be independent of each other. Dominance behaviour is represented as high rates of approaching and displacing on PC1 and as the opposing loadings of David’s score (indication of dominant status) and submissions (indication of subordinate status) on PC5. Dominant juvenile *A. burtoni* approach and displace at significantly higher rates than subordinates (Solomon-Lane et al., 2022). For PC5, being in or near the territory zone, with less time in the investigate zone, was associated with low status when given the opportunity to be dominant. This is mirrored on PC1 by the negative association between being in the territory zone and approaching and displacing in the dominance behaviour assay. The relative dominance of juveniles from the socialized pairs is also consistent with having significantly lower directional consistency (i.e., more agonistically symmetrical) in the subordinate behaviour assay. In our previous study, group-reared juveniles were less submissive in a subordinate role than pair-reared juveniles (Solomon-Lane & Hofmann, 2019). It is possible this was not the case for stable group juveniles in this study due to group size differences (6 fish vs. 16 fish).

The most striking difference for PC5 is the involvement of cortisol. We found no treatment differences in water-borne cortisol levels, and PC5 was the only PC on which cortisol loaded strongly. Higher cortisol was associated with lower social status, less exploratory behaviour, and more time in the territory away from the novel cue fish. This suite of behaviours with cortisol resembles previously identified syndromes (Réale et al., 2007), such as the pace-of-life syndrome (Careau & Garland, 2012) and coping styles (Koolhaas et al., 1999). A “fast” pace-of-life is associated with increased activity, exploration, boldness, and aggressiveness, along with one or more traits from the slow-fast metabolic continuum (Careau & Garland, 2012). Coping styles are an integrative phenotype in which a behavioural syndrome aligns with stress physiology (Koolhaas et al., 1999). Coping styles have been observed across species (Alfonso et al., 2019; Conrad et al., 2011; Øverli et al., 2007) and are sensitive to early-life effects (Sih, 2011). Behaviourally, proactive copers are more active, aggressive, and bold compared to reactive copers. In response to stress, proactive copers have higher sympathetic reactivity and lower HPA/I activity (Koolhaas et al., 1999). Whether socialized pairs (this study) or group-reared juveniles (Solomon-Lane & Hofmann, 2019) exhibit a fast-pace-life or are proactive copers are hypotheses that should be tested directly. The association between high cortisol and low status is consistent with previous studies of juvenile *A. burtoni*, which showed higher whole brain GR1a and GR1b expression, and lower GR2 and MR expression, in fish with higher dominance scores (Solomon-Lane et al., 2022). Efficient negative feedback may be mediated, in part, by GR1 expression, leading to lower cortisol levels (Solomon-Lane & Hofmann 2019). The relationship between adult *A. burtoni* status and cortisol varies across studies and can be elevated in subordinates (Maruska et al., 2022). Adult *A. burtoni* do not appear to form coping styles (Butler et al., 2018), and this suite of behaviour is not always correlated with stress responsiveness across species (e.g., Thomson et al., 2011). Overall, early-life exposure to social complexity tends to benefit social competence and skills, whereas social instability tends to have lasting, negative phenotypic effects, such as elevated HPA/I axis activity, weight loss, elevated aggression, and decreased activity (Kohn, 2019). This suggests that exchanging members of the pair in the socialized treatment may not have been perceived as instability, potentially because of the predictable schedule (Kohn, 2019). An alternative explanation is that familiarity played a role in the treatment differences between socialized pairs vs. stable groups and pairs. The dominant and subordinate assays were highly similar to the way we socialized the pairs, and familiarity with the assay could lead to appearing more active, interactive, and dominant. Future studies can test these potential behavioural mechanisms directly.

We observed fish in their rearing environments to gain insights into the specific social experiences and social sensory cues, or proximate behavioural mechanisms, responsible for early-life social effects (Taborsky, 2016). Unsurprisingly rates of behaviour were higher in the stable groups compared to the pairs. Rates of behaviour per fish did not differ across treatment groups, and there were no differences in total agonistic efficiency. These data confirm that a group (compared to pair) social environment has more opportunities for social experiences, such as direct involvement in an interaction and observations of others interacting (Desjardins et al., 2012; Oliveira et al., 1998). The mean distance between dyads was also significantly larger in stable groups than the pairs, and although the tank was larger for groups, the smallest dyad distance was significantly smaller in groups than social pairs. There were no differences across treatments when scaled for the size of the tank. We were unable to identify and track individuals because the ink tags were not visible on video. As a result, we could not quantify individual social experience, social status, social network position, or spatial position, which we hypothesize are causally related to behavioural development and phenotype (Kasumovic, 2013; Taborsky, 2016). The evidence that social interactions are assortative within groups is overwhelming across species (e.g., Chase et al., 2022; Croft et al., 2005, 2009; Pike et al., 2008; Williamson et al., 2016), and juvenile *A. burtoni* form social status relationships and nuanced social structures (Solomon-Lane et al., 2022). Therefore, we expect social status experience and the degree of agonistic asymmetry across social partners to be particularly important for behavioural development. In addition to its role as an indicator of social relationships, spatial proximity may also affect social sensory cue perception and communication, for example, via mechanosensory and chemosensory cues (e.g., in adult *A. burtoni*, Butler & Maruska, 2016; Nikonov et al., 2017) that could be stronger for fish in closer proximity.

Overall, our work demonstrates that multiple behavioural mechanisms—the number of early-life social partners and social stability or familiarity—affect juvenile *A. burtoni* development and phenotype, including integrated behavioural and neuroendocrine traits. In addition to manipulating social experience directly, we observed juveniles in their rearing environments, which are necessary steps towards understanding the social experiences accrued during development and the mechanisms by which early experiences exert long-term and disproportionately strong effects on adult phenotype (Buist et al., 2013; Jonsson & Jonsson, 2014; Kasumovic, 2013; Taborsky, 2016, 2017). Although the simplistic social contexts we used in this study are unlikely to reflect the dynamics of f larger, more complex groups found in nature (Chase et al., 2003), we expect that these behavioural mechanisms, and potentially others—acting additively or synergistically—will also be influential in more naturalistic contexts. Testing the hypotheses we generated here will be key to uncovering the behavioural mechanisms of developmental plasticity and phenotype, as well as the role of neuroendocrine stress axis function, in *A. burtoni* and other social species.

## ACKNOWLEDGEMENTS

This work was supported by start-up funds from the W.M. Keck Science Department to TKSL, W.M. Keck Science Department Summer Fellowships to DDB, HLG, IPH, and EAM; Claremont McKenna College Sponsored Internships & Experiences to JL, and the Claremont Colleges Intercollegiate Neuroscience Summer Research Fellowship to JW. We thank Maya Shah for contributing to scoring behaviour.

## DATA AVAILABILITY

Data will be made available in an online repository (Dryad).

