## Supplemental Info for "Multiple behavioural mechanisms shape development in a highly social cichlid fish"

**Supplementary Information for: Multiple behavioral mechanisms shape social behavioral development in a highly social cichlid fish**

**Supplemental Table 1:** Standard lengths (SL, mm) of fry from the 7 broods from which experimental individuals were selected.

| Brood ID | Brood size | Mean SL | SEM | Median SL | Min SL | Max SL |
|---|---|---|---|---|---|---|
| Brood 3 | 15 | 5.26 | 0.102 | 5.17 | 4.68 | 6.06 |
| Brood 4 | 21 | 5.20 | 0.0696 | 5.12 | 4.59 | 5.81 |
| Brood 5 | 18 | 6.19 | 0.143 | 6.17 | 4.70 | 7.31 |
| Brood 6 | 19 | 7.16 | 0.145 | 7.17 | 5.19 | 8.13 |
| Brood 7 | 6 | 5.86 | 0.114 | 5.75 | 5.63 | 6.36 |
| Brood 8 | 4 | 6.13 | 0.0675 | 6.16 | 5.94 | 6.26 |
| Brood 9 | 4 | 5.83 | 0.287 | 5.73 | 5.26 | 6.62 |

Brood size is the number of fry removed from a single mother's buccal cavity. Standard length (SL) was measured using ImageJ (Schneider et al., 2012) from digital images of the fish next to a ruler.

**Supplemental Table 2:** Treatment differences in home tank social behavior and distances.

| Behavior / measure | DF | Test Statistic | p-value |
|---|---|---|---|
| **Approaches per fish** | 2, 38 | F=0.86 | 0.43 |
| **Displacements per fish** | 2, 37 | F=1.83 | 0.18 |
| **Into territory per fish** | 2 | $\chi^2$=1.43 | 0.50 |
| **Scaled mean dyad distance** | 2, 46 | F=1.1 | 0.34 |

Results of one-way ANOVAs (F) or nonparametric Kruskal-Wallis tests ($\chi^2$). Eta-squared is reported for effect sizes. Significant results in bold. Tukey HSD tests were used for *post hoc* analysis of significant ANOVA results. Dunn's tests were used for *post hoc* analysis of significant Kruskal-Wallis results.

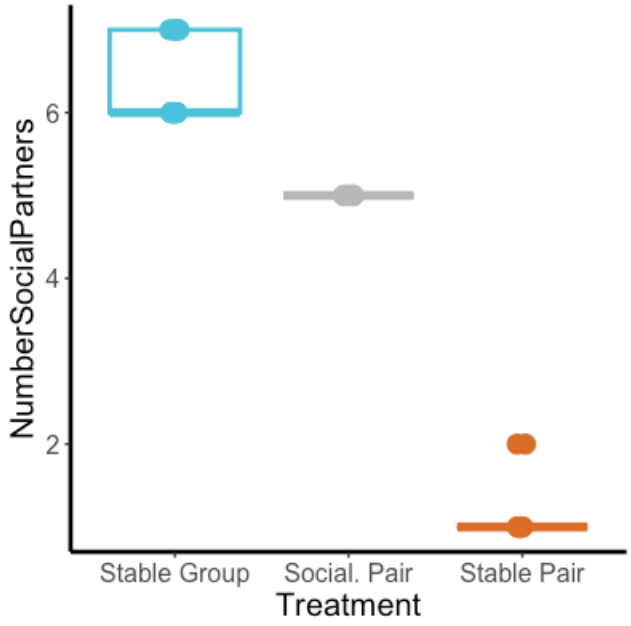


**Supplemental Figure 1:** The number of social partners experienced by fish in each home tank and treatment. Socialized pairs had exactly 5 partners. The stable pairs with 2 social partners and stable groups with 7 social partners had a fish die and was replaced.

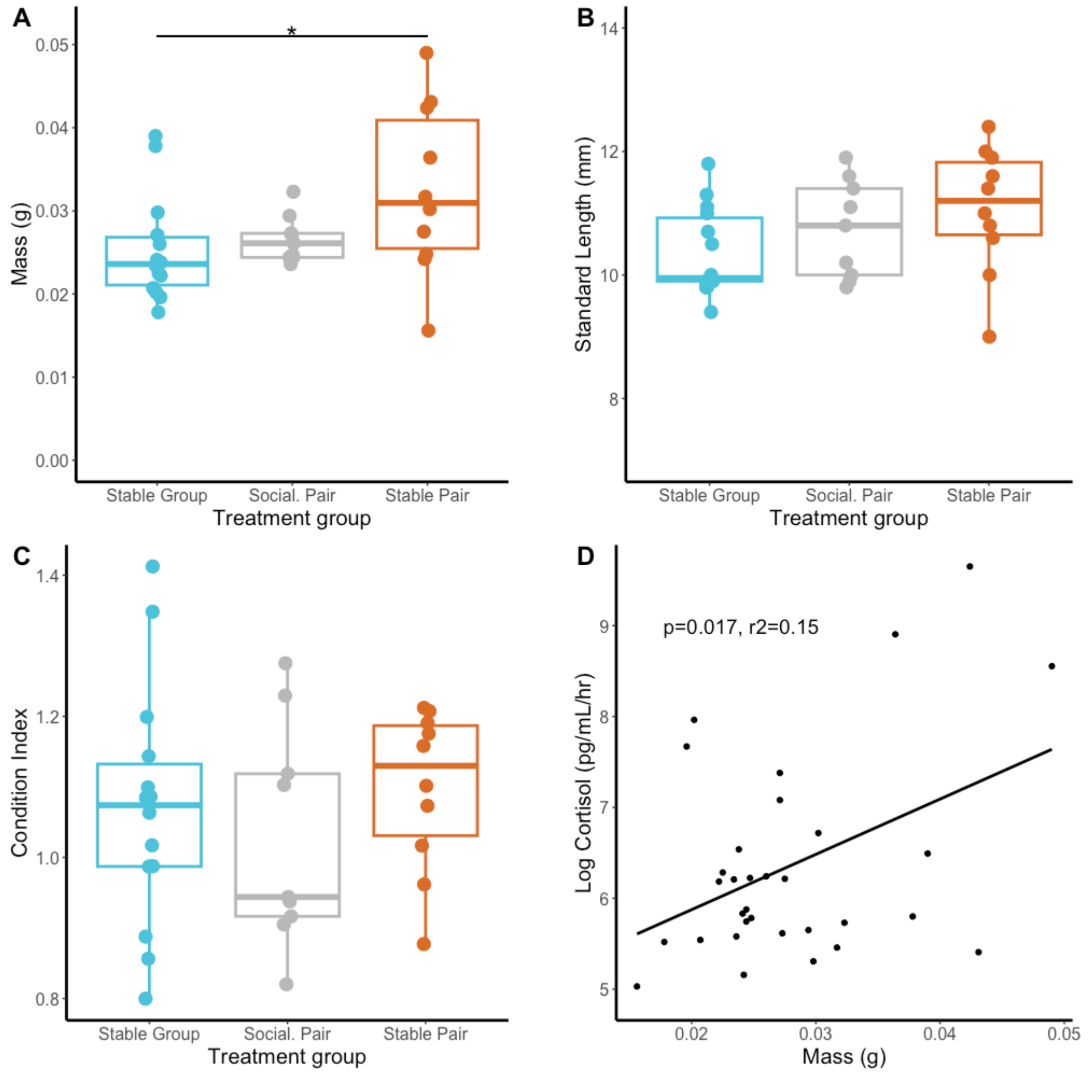


**Supplemental Figure 2:** A) Mass (g) and B) standard length (mm) of juveniles measured at the end of the experiment, on the day of individual behavior testing. C) Condition index was calculated as Fulton's *K*. D) Association between mass and water-borne cortisol levels. *p<0.05

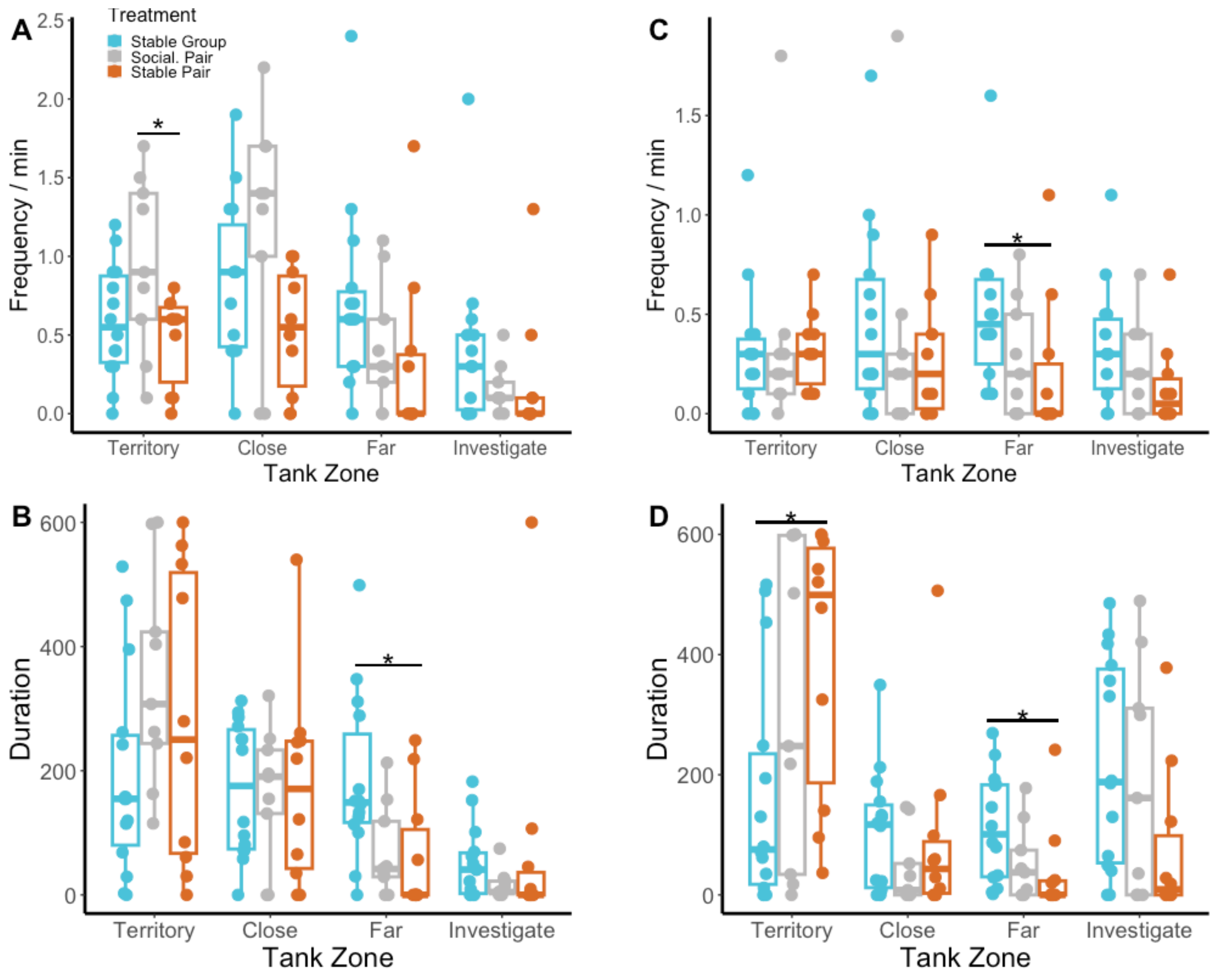


**Supplemental Figure 3:** In the open field exploration, A) the frequency of entering and B) the total time (s) spent in each zone of the tank. In the social cue investigation, C) the frequency of entering and D) the total time (s) spent in each zone of the tank. *p<0.05.

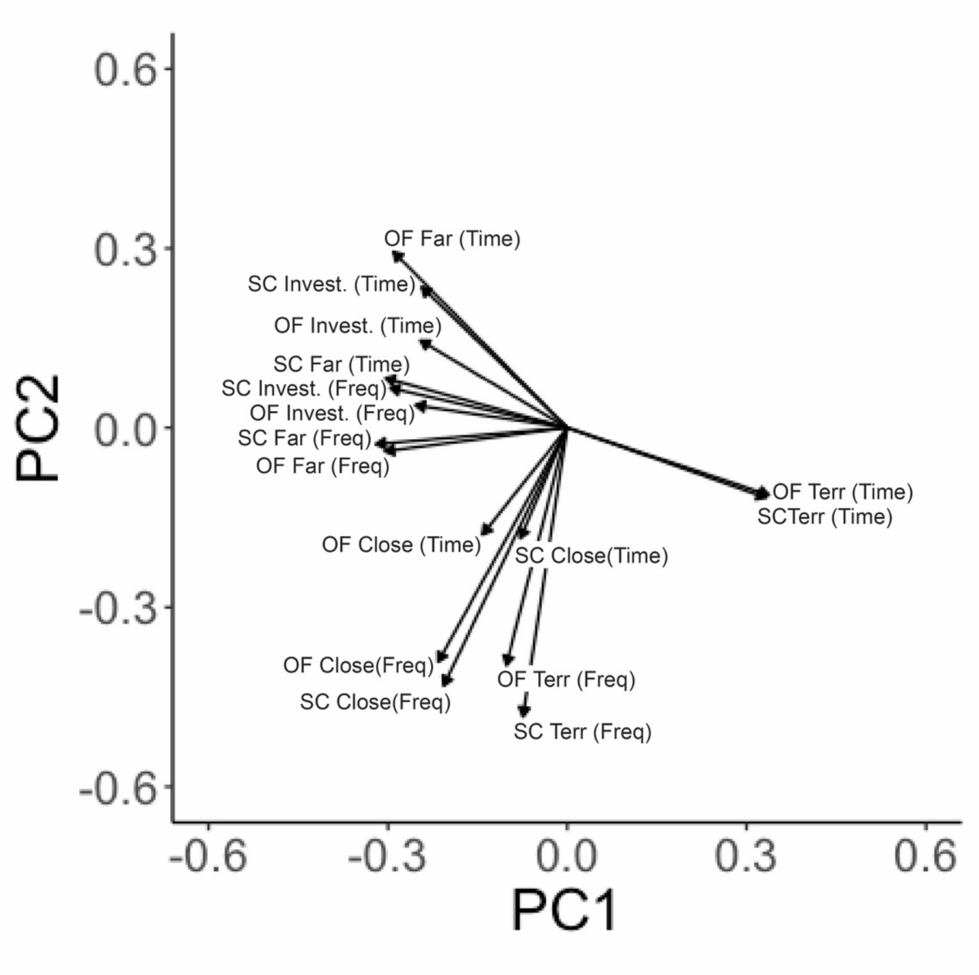


**Supplemental Figure 4:** Principal components analysis (PCA) of open field exploration (OF) and social cue investigation (SC) frequency entering and time in each zone of the tank: investigate (invest.), close, far, and territory (terr) zones. The vector plot shows the PC variables that load on PC1 and PC2. Because of the strong concordance between OF and SC vectors for a given zone and measure (time, frequency), these data were used to determine which variables to include in the larger PCA with variables from all four behavior assays. All SC variables were included in the larger PCA. Only OF variables that did not closely align with the corresponding SC variable were included: OF time in the far zone and OF time in the investigate zone.

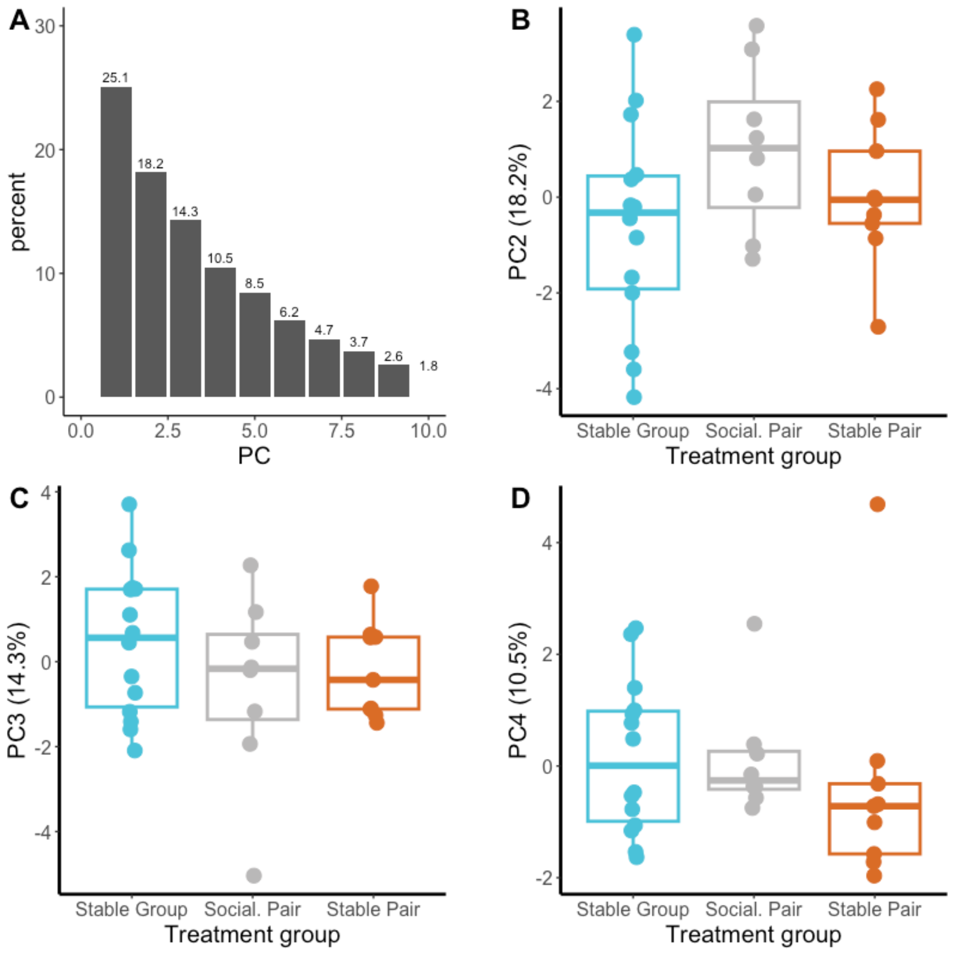


**Supplemental Figure 5:** Principal components analysis (PCA) of cortisol and behavior—including open field time in the far and investigate zones; social cue time in and frequency entering the territory, close, far, and investigate zones; and dominant and subordinate behavior assay approaches, displacements, approaches received, submissions, and David's Score. A) Skree plot of the percentages of the variation in the data explained by each PC. B-D) Treatment differences in PC2-PC4.
